# Layer-specific stimulations of parvalbumin-positive cortical interneurons in mice entrain brain rhythms to different frequencies

**DOI:** 10.1101/2021.03.31.437894

**Authors:** François David, Mélodie Borel, Suleman Ayub, Patrick Ruther, Luc J. Gentet

## Abstract

Neocortical interneurons provide inhibition responsible for organizing neuronal activity into brain oscillations that subserve cognitive functions such as memory, attention or prediction. However, little is known about the interneuronal contribution to the entrainment of neocortical oscillations within and across different cortical layers. Here, using layer-specific optogenetic stimulations with micro-Light-Emitting Diode (µLED) arrays, directed toward parvalbumin-expressing (PV) interneurons in non-anesthetized awake mice, we found that supragranular layer stimulations of PV neurons were most efficient at entraining supragranular local field potential (LFP) oscillations at gamma frequencies (γ: 25 - 80 Hz), whereas infragranular layer stimulation of PV neurons better entrained the LFP at delta (δ: 2 - 5 Hz) and theta (θ: 6 - 10 Hz) frequencies.

At the level of neuronal action potential activity, we observed that supragranular neurons better followed the imposed PV stimulation rhythm than their infragranular counterparts at most frequencies when the stimulation was delivered in their respective layer. Moreover, the neuronal entrainment evoked by local stimulation could propagate across layers, though with a lesser impact when the stimulation occurs in deep layers, suggesting an orientation-selective propagation. These results establish a layer-based framework for oscillation to entrain the primary somatosensory cortex in awake conditions.

## Introduction

Neocortical brain rhythms are emergent neuronal circuit properties that underlie many functional mechanisms, such as plasticity (Chauvette et al., 2012; Durkin et al., 2017; Peyrache and Seibt, 2020), synaptic integration (Wehr and Zador, 2003) and inter-area communication (Fries, 2015). Locally, they manifest themselves as electrophysiological bouts of oscillations at frequencies from 0.05 Hz to 500 Hz, visible at the EEG level. Their emergence and maintenance both depend on tight interactions between excitatory and inhibitory inputs (Tiesinga and Sejnowski, 2009), as well as intrinsic neuronal properties (Wang, 2010).

Among the different subclasses of cortical GABAergic interneurons, parvalbumin-containing (PV) fast-spiking (FS) neurons, mostly basket cells have been shown to play a critical role in the emergence, maintenance and termination of sleep slow waves (Shu et al., 2003; Zucca et al., 2019) and in the pacemaker activity of γ oscillations (Cardin et al., 2009; Sohal et al., 2009; Kuki et al., 2012; Welle and Contreras, 2016) through the strong and rapid inhibition they provide onto the soma and proximal dendrites of pyramidal neurons (Gentet et al., 2012; Tremblay et al., 2016). Indeed, their neuronal activity has been shown to be correlated with various brain rhythms throughout the cortex (Klausberger et al., 2005; Peyrache and Destexhe, 2019). Therefore, cortical FS cells, which mainly consist of GABAergic PV interneurons (Hu et al., 2014) play a major role in controlling the excitability of the network (Ferguson and Gao, 2018) through their widespread connections to local pyramidal neurons (Packer and Yuste, 2011).

Brain oscillations modulate both bottom-up and top-down cognitive processes (Arnal and Giraud, 2012; van Kerkoerle et al., 2014; Bastos et al., 2015, 2020) through the layer-based functional architecture of the neocortex. Sensory inputs mainly enter the cortical column at the level of granular layer 4, are then transferred to layer 2-3, which project to layer 5-6 (Shipp, 2007). Contextual inputs are carried into supragranular layers 1-3 (Douglas and Martin, 2004; Phillips et al., 2016), while feedback input from higher cortical areas control the input-output function of the column at multiple levels of the infragranular layers 5-6 (Ahissar and Staiger, 2010) and supragranular layers (Shipp, 2007). Given the importance for cognitive processes of rhythmic dynamics and the layer-based neocortical architecture, we hypothesized that the local excitatory-inhibitory interactions responsible for oscillations would selectively impact those rhythms based on their location, thus differentially impacting frequency amplification across neocortical layers. How the various PV cells that are present in supragranular, granular and infragranular layers entrain and transfer different brain rhythms within a cortical column remains to be determined.

To test this hypothesis, we investigated the differential contribution of infragranular, granular and supragranular layer PV interneurons to the amplification of the oscillatory regimes of the mouse primary somatosensory barrel cortex (S1) via rhythmic and local activation of local PV-containing interneurons with arrays of miniaturized sources of light (µLEDs – see Ayub et al., 2017, 2020). In non-anaesthetized head-fixed mice during quiet wakefulness in which spontaneous activity covers a large range of spontaneous oscillation frequencies (0.5 - 100 Hz, Buzsáki and Draguhn, 2004), and using a combination of virally-mediated cre-dependent channelrhodopsin expression in transgenic PV-cre mice (Madisen et al., 2012) combined with neuronal ensemble recordings, we optogenetically activated PV neurons located in either supra- or infragranular layers of S1 and observed their impact at the level of local field potentials (LFP) and cortical neuronal activity in all layers. We uncovered stronger impact of infragranular-layer PV activation at δ and θ frequencies on LFPs throughout the column, while supragranular PV cell activation better entrained oscillations at γ rhythms (25 - 80 Hz). Furthermore, we found that supragranular cortical neurons were better entrained than their infragranular counterparts at γ frequencies and in the low frequency range i.e. slow wave and δ frequencies. PV mediated inhibition evoked in supragranular layers more strongly impacts neuronal activity in rest of the neocortical column than infragranular layer evoked inhibition impacts upper layers. Our results reveal how cortical PV neurons can differentially impact neuronal and network neocortical rhythms depending on their anatomical location in various layers.

## Material and Methods

### Transgenic mice

All animal experiments were conducted after approval by the local ethical committee of the University of Lyon and the Ministry of Science (Protocol number: Apafis #4613) in accordance with the European directive 86/609/EEC on the protection of animals used for experimental and other scientific purposes. We used offsprings of PV-Cre driver mice (008069, Jackson). All animals used in this study were group-housed in the vivarium under normal light cycle conditions.

### Surgery and recording

All surgical procedures were performed under 1.5-2% isoflurane anaesthesia, after prior injection of 5 mg/kg carprofen. First, animals (6-12 wk old) were implanted with a lightweight head-bar and a recording chamber. A small craniotomy was performed 300 μm lateral to the C2 barrel column (identified through intrinsic optical imaging) and adeno-associated virus (AAV1.EF1a.DIO.hChR2(H134R)-eYFP.WPRE.hGH; Penn Vector Core, University of Pennsylvania) was injected into the C2 barrel column at depths of 250 μm, 500 μm, 750 μm and 1000 μm (100 nl per injection site). Mice were then progressively habituated to head-fixed conditions with regular rewards for 10 days before recording. On the day of recording, the craniotomy at the site of viral injection was enlarged, a second small craniotomy was performed above the C2 column and the dura was removed at both locations 300μm apart in anticipation for the recording electrode and µLED array insertions. Animals were allowed to recover from anaesthesia for at least 2 h, during which the craniotomy was kept humidified with artificial cerebrospinal fluid (ACSF). The µLED array and a linear 32-channel silicon electrode were slowly lowered into the brain, with the light sources from the µLED array facing the C2 barrel column (Figure 1a). The electrode and the µLED array were fixed together with a small amount of silicon gel (Kwik-Cast) for higher mechanical stability. Signals were acquired full-band and continuously using a Multi-Electrode-Array portable 32-channel (ME32-FAI-mPA, Multichannel Systems MCS GmbH) at 20 kHz. Optogenetic stimulation time points were simultaneously recorded on an extra-digital channel.

**Figure 1:**
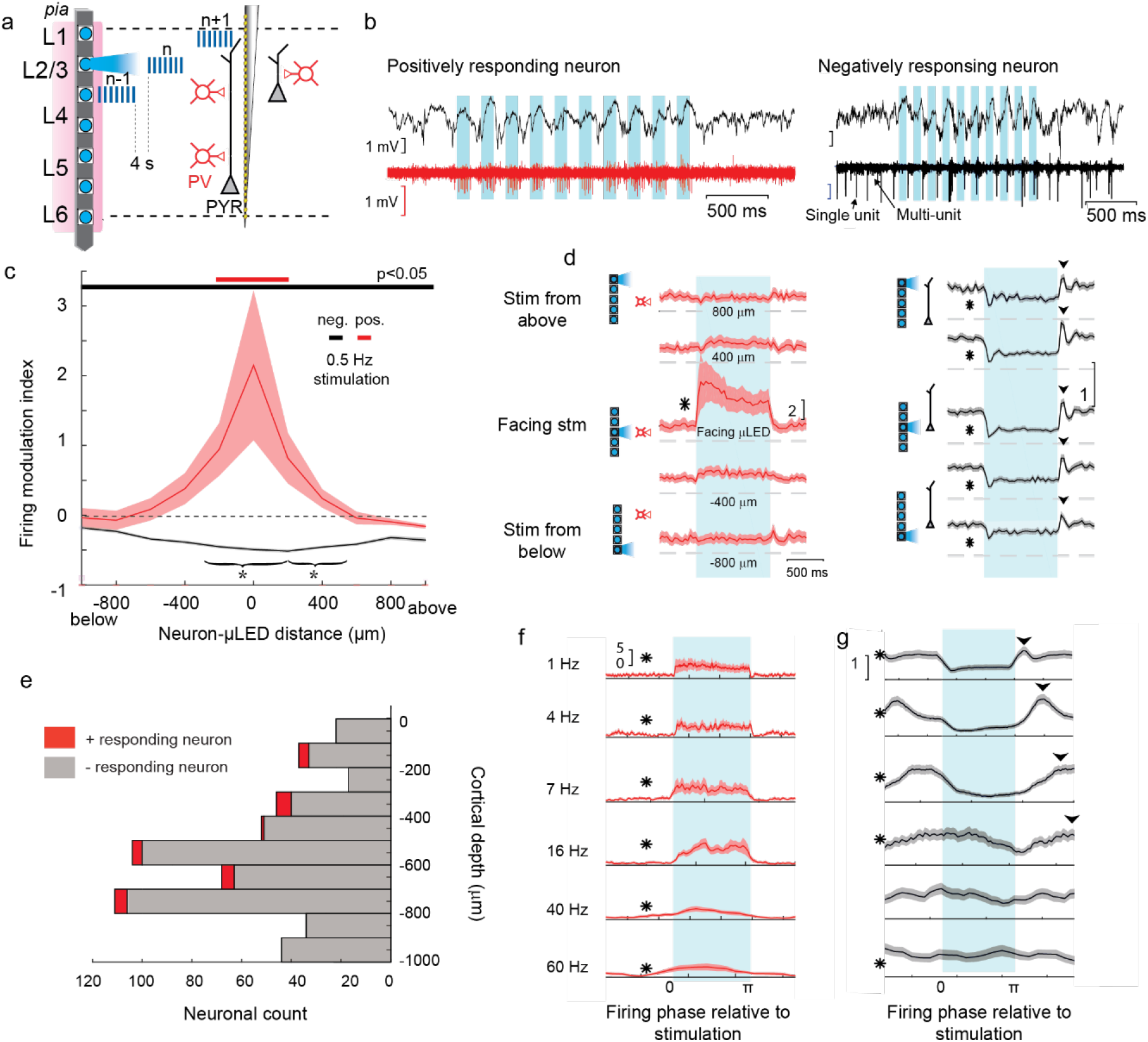
Parvalbumin-positive neurons control of inhibition within a cortical S1 barrel column. **a**) Schematic positioning of the µLED array and multi-site electrode facing each other across cortical layers (L1-6) of a mouse S1 barrel column. PY: pyramidal neurons, PV: parvalbumin positive interneurons. µLED light pulses were sequentially emitted every 4 seconds. **b**) Examples of LFP (top) and high-passed filtered traces for positively (red) and negatively (black) responding units. Light-ON periods of stimulation are indicated in shaded blue. **c**) Average neuronal firing modulation (± SEM) at various distances from the facing µLED (negative values on the x-axis indicate responses of neuron below the µLED) for negatively (neg) and positively (pos) responding population of neurons at a stimulation frequency of 0.5 Hz. Horizontal color-coded bars indicate values significantly different from 0 (p < 0.05, Wilcoxon rank sum test). * indicate the distance from which the inhibition starts being significantly different from the maximal inhibition. **d**) Average normalized response at 0.5 Hz stimulation (± SEM (shaded area)) of positively-responding (left) and negatively-responding neurons based on neuronal location relative to the µLED ‘ON’ (blue rectangle). * indicates p-value < 0.05, Wilcoxon signed-rank test for differences relative to µLED ‘OFF‘ times. Arrowheads indicate the post-inhibition rebound peak of activity. **e**) Depths distribution of all responding neuronal units based on their category (n = 7 mice). **f**) Average response of positively-responding neurons for various light pulse train frequencies delivered from the facing µLED. Time basis is adapted to the period of stimulation. **g**) Same as f) for negatively-responding neurons. Note the shift on the x-axis (phase) of the post-inhibitory rebound (arrowheads) for frequencies below 16 Hz.

### Optogenetic stimulation

We used two sets of µLED array types (Ayub et al., 2020) that differed only by their inter-µLED distances (µLED pitch values of 150 or 250µm). The wavelength of blue light peaked at 455 nm. Individual µLEDs were controlled using a custom-made Arduino-based driver. Light intensity was set to 3 µW (E =0.4 mW.mm^−2^) which was previously found to be sufficient to activate µLED-specific subsets of constitutively-expressing ChR2 PV neurons on the densest array (Ayub et al., 2020) while spatially restricting the light-activated neurons. Trains of 10 light pulses interspaced by 4 seconds (Figure 1a) mimicked the transient aspects of brain oscillations which often occur in short bouts. The trains consisted of Light-On and Light-Off sequences each pulse lasting half of the stimulation period. This 50% ratio of Light-On times was kept constant across stimulation frequencies (for example for 10 Hz stimulations, light was on 50 ms, then off for 50 ms and so on). The stimulation frequencies were chosen to cover a wide range of physiological brain rhythms (i.e. 0.5-100 Hz: Slow waves (0.5-1 Hz), Delta (δ, 2-5 Hz), Theta (θ, 6-10 Hz), Beta (β, 20-25Hz) and Gamma (γ, 25-80 Hz)). Each stimulation frequency-LED combination was repeated 6 to 10 times per experiment in a randomized order.

### Sensory stimulation and C2 barrel identification

On the day of experiments, whiskers were trimmed except for the C2 whisker, which was subsequently coated with ferromagnetic paint. An electromagnetic coil (Gotronics, 24 Vcc MS5030-24) was positioned at a distance of ∼1cm in front of the C2 whisker. A 5ms electromagnetic pulse evoked a deflection of the whisker which could be tracked under IR light.

### Analysis

The local field potential (LFP) was derived from the raw signal downsampled at 1 kHz. For current source density (CSD) analysis (Mitzdorf, 1985), the transforms were computed locked on the light pulse stimulations or on the magnetic stimulations. Values were interpolated and averaged over a 160 μm sliding window. The LFP was bandpass-filtered around the stimulation frequency (±20%) for the CSD analysis. Depth landmarks during PV-stimulation could be identified and positioned for all mice based on the limits between sinks and sources transitions across cortical depths. Those did not depend on the position of the glowing μLED. Only granular layer (layer 4) positions were used and delimited by using the CSD performed following sensory stimulations (Swadlow et al., 2002) and layer 5 position (Figure 2). The granular inferior limit was estimated based on the gradient-descent clustering process that generated 3 clusters grouped on the basis of channel pair γ(25-80Hz)-coherence (Berényi et al., 2014) performed on short samples of spontaneous LFP activity. This limit was consistently found to be above the peak of MUA neuronal firing (Senzai et al., 2019). Then, we defined the position of the granular layer superior limit, 150 μm above the inferior limit according to the accepted granular layer thickness (Bopp et al., 2017). Neurons could then be assigned to either the supragranular, granular or infragranular layer.

**Figure 2:**
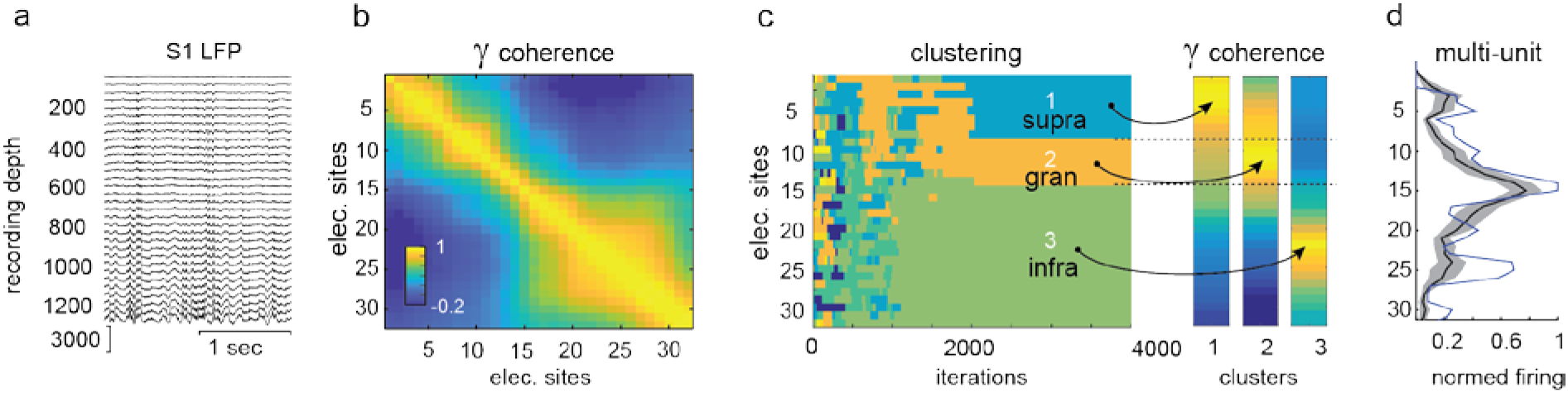
Electro-anatomical characterization of neocortical layers. **a)** LFP recording example along laminar electrodes positioned in S1 barrel cortex. **b**) LFP coherence (color coded) in the gamma (γ) frequency band between each recording site pairs. **c**) γ-coherence-based clustering of recording sites (left). Coherence of 3 recording sites from each found clusters (right). Dashed lines indicates the borders of the clusters. **d**) Total firing rate per cortical depth of recording for the same mouse as in (a) in blue normalized to 1 at its peak value. Average (black) ±SEM (shaded grey) for all mice (n=7).

For LFP analysis, a recording site within the granular layer was used as a common local reference site for supra- and infra-granular differential LFP measurements (Herreras, 2016) and therefore subtracted from the infra and supra LFP recorded traces. To measure the changes evoked by each stimulation frequency, we performed a frequency analysis on the corresponding LFPs using short-time Fourier transforms. The LFP modulation index was the ratio of this power measure and a control value obtained from the concatenated signal outside the stimulation intervals for a number of samples identical to the number of trials randomly pooled. This control value was then computed 60 times to avoid sample size variability during the control period and then averaged to get the control value. The LFP modulation index was then derived from the relative change of power during the light stimulation compared to the one outside the stimulation (an index of 0 indicates no change and 1 indicates a 2-fold increase).

Single- and multi-unit neuronal activity was extracted from the high-pass filtered signal, automatically, clustered using Klusta suite software (https://klusta.readthedocs.io/, (Rossant et al., 2016)) and manually curated. Clusters were merged only when multiple evidences were found indicating that the spikes likely belong to the same neurons, based on spike shape, channel distribution of spike shapes and temporal evolution (likely due to the electrode spatial drift). The rare manual interventions explain the high number of clusters as the clustering algorithm settings tend to overestimate their number. Artifact-contaminated neuronal clusters were manually excluded based on waveform and autocorrelograms. Neuronal clusters were then categorized either as belonging to putative inhibitory (INH) neurons (mostly fast-spiking (FS) neurons) if their spike half-width durations were less than 0.6 - 0.75 times their trough-to-peak durations or to putative regular-spiking (RS) excitatory (EXC) neurons in all other cases (Barthó et al., 2004). In vitro and in vivo recordings indicate that inhibitory fast-spiker neurons have a narrower spike shape (McCormick et al., 1985; Mitchell et al., 2007; Tseng and Han, 2021). 74% of the neurons that responded to optogenetic stimulation had a narrow spike shape (Figure 1d), others could be PV+ neurons that are not fast-spiker neurons (Hu et al., 2014). We will therefore refer to these neuronal classes based on cluster spike shape as ‘putative’ INH and EXC neurons. Clusters with a smaller firing rate than 0.1Hz were excluded from further analysis, as the analysis is based on relative change of firing and low firing rate induce less reliable baseline. Units were further categorized with respect to their response to µLED stimulation into ‘positively responding’ if their firing rates more than doubled (modulation index greater than 1) or ‘negatively responding’ if the firing rates simply decreased. Units that did not fall within these criteria were discarded from the analysis. A neuronal firing modulation index was defined as the average difference between the firing rate observed during the light stimulation and the firing rate measured in-between light pulses, divided by this latter. The modulation index quantifies how the neuronal population fire relative to the stimulation. The neuronal coherence was estimated as the module of the spike phase histograms relative to the optogenetic stimulation. The neuronal coherence quantifies how individual neurons consistently fire relative to the stimulation. The neuronal angle was estimated as the average phase of firing of neurons. Phase 0 is the start of the μLED pulse and π is the start of the gap in between the pulses. For representation purposes, curves were smoothed using a 3 points window moving average. Neuronal clusters recorded below 1000μm potentially did not belong to the cortex but rather to corpus callosum. While we recorded neuronal clusters below 1000 μm, The <u>Allen</u> <u>Institute Brain Atlas</u> indicates that mouse cortical thickness can reach in excess of 1 mm. Results were not affected by the inclusion or exclusion of these recorded neurons.

### Statistics

Non-parametric paired or unpaired (Wilcoxon signed or unsigned, respectively) tests were performed on modulation indexes. Phase distributions were tested for non-uniformity with the Rayleigh test of uniformity of circular data and phase distribution pairs were compared with the Watson-William test for homogeneity of the means. The distribution modality of modulation index relative to stimulation frequency was assessed using a Hartigan dip test (Hartigan and Hartigan, 1985). We reported significance when p-values < 0.05. Results are reported as means ± standard errors of the mean (SEM), which is extended in the circular domain by using the circular mean and the circular deviation over the square root of the sample size.

## Results

We first explored how a rhythmic and specific activation of a pool of local PV neurons impacted neuronal activity in the mouse primary somatosensory cortex (S1) based on depth and frequency (Figure 1a-b, Supplementary Figure 1a-b). Only neurons with a baseline firing rate above 0.1 Hz were considered in order to better measure relative changes induced by the optogenetic stimulation of PV neurons. 60 neurons out of a total of 669 clusters (n=7 mice) were therefore discarded.

### Local and global inhibition induced by rhythmic activation of PV neurons in the mouse primary somatosensory cortex

Spike shape analysis of selected neuronal clusters (n=609) indicated that 20 out of 27 (74%) positively-responding neurons (example Figure 1b left) were putative inhibitory based on their action potential waveforms (Supplementary Figure 1c-e), and were therefore categorized as positively responding PV cells. Importantly, maximal neuronal response was always observed when the positively-responding neuron was facing the stimulated μLED (Figure 1c,d left) while all categories of recorded neurons were distributed along the cortical depth (Figure 1e). Therefore PV-neuronal activation was spatially-constrained to cortical layers in the vicinity of the stimulated μLED, confirming our previous results (Ayub et al., 2020).

The remaining population of recorded neurons displayed either decreased firing (Figure 1b-d right, n = 548 neurons) or failed to reach criteria for a change detection in their spiking rate in response to light pulses (n = 34 neurons). In contrast to the positively-responding neurons, the decrease in firing of the negatively-responding neurons spread well beyond the cortical depth facing the stimulating µLED (Figure 1c).

Amongst the inhibited neuronal clusters, a majority (370 out of 548) was classified as putative excitatory neurons (EXC. neur) based on their broad spike-shape typical of pyramidal regular-spiking neurons (RS) (Supplementary Figure 1c-f). Interestingly, a large number of narrow-spike shaped putative inhibitory (INH) neurons, also responded with decreased firing rates (121 out of 148 neuronal clusters, Supplementary Figure 1d-e).

While positive neuronal responses to μLED-stimulations were observed across most stimulation frequencies for positively responding neurons (Figure 1f, Supplementary Figure 1g), decreased neuronal responses were not observed across all frequencies (Figure 1g, Supplementary Figure 1h). In order to explore systematically the spatial and temporal effect of stimulation, we tested how light stimulations at frequencies spanning from 0.5 to 100 Hz differentially modulated neuronal activity across the cortical column (Figure 1f-g, Supplementary Figure 1i). While positively-responding neurons remained rhythmically entrained at nearly all stimulation frequencies, their modulation appeared reduced (and insignificant) at some high frequencies (for instance at 100Hz, Supplementary Figure 1i). Similarly, the negative modulation weakened with stimulation frequencies above 7 Hz (for example at 40Hz; Figure 1g, Supplementary Figure 1i). Together, these results reveal the existence of a cut-off frequency in the ability of local PV neurons to rhythmically entrain the overall activity of the neocortical network at higher stimulation frequencies.

Moreover, stimulation frequency further affected the spatial extent of neuronal modulation. Responses from positively-responding neurons were spatially constrained at very low frequencies (0.5 Hz, red curve in Figure 1c; Facing μLED modulation index = 2.15 ± 1.07, n = 18, p = 0.003; 400 µm apart: modulation index = 0.38 ± 0.22, n = 14, p = 0.11; non-zero mean, Wilcoxon rank sum test). On the other hand, the firing rates of positively-responding neurons up to 600 µm apart, was still significantly increased at some high (60 Hz, yellow curve, Supplementary Figure 1j; Facing μLED: modulation index = 1.48 ± 0.42, n = 16, p = 0.0009; −600 µm apart: modulation index = 0.43 ± 0.16, n = 13, p = 0.005; non-zero mean Wilcoxon rank sum test).

In contrast, the rhythmic inhibition remained significant at low frequencies for a large depth range for example at 0.5 Hz (Figure 1c), while being maximal when light was delivered from nearby µLEDs (+200μm) (Figure 1c; modulation index = −0.51 ± 0.02, vs. 400 µm apart, modulation index = −0.53 ± 0.02, p = 0.0003). The phasic inhibition became weaker at higher frequencies although still significant (Supplementary figure 1j, modulation index at 60 Hz = −0.21 ± 0.011, p =0.002) and lacking layer-specificity (Supplementary figure 1j). This suggests that higher frequency stimulations are more prone to equal spreading of rhythmic inhibition throughout layers, albeit at a weaker intensity.

### Layer-specific neuronal entrainment by PV rhythmic neuronal activation

Since PV-driven circuit inhibition has stronger effects locally than further away within a cortical column, we next wanted to know whether different cortical layer circuits were more prone to the amplification and tuning of specific neocortical rhythms. To compare the entrainment properties between layers, we first needed to determine some electro-anatomical boundaries (Berényi et al., 2014), to define those layers across stimulation frequencies. We used LFP spontaneous activity (Figure 2a) and γ-coherence estimated across pairs of recording sites (Figure 2b) to cluster the channels with a gradient-descent algorithm (Berényi et al., 2014). This yielded the separation of 3 compartments (Figure 2c) whose boundaries matched the anatomically-defined infragranular, granular and supragranular cortical layers (Berényi et al., 2014). The average depth of the infragranular-granular boundary was −505±42 µm (n=7) whereas the average depth of the granular-supragranular layer boundary was 277±45 µm (n=7). In addition the total neuronal firing rate was estimated along the recording sites of the electrode (Figure 2d) for each experiment. The position of maximal firing, previously associated to layer 5 position (Senzai et al., 2019) was found at - 594±18 µm (n=7) just below the granular layer inferior limit. As those relative positions were consistent across mice, we chose those references to define this position as the infragranular and granular layer limit positions. We used a standard granular layer thickness 150um (Bopp et al., 2017) to define this position relative to the infragranular layers. Lastly to cross-validate those layer positioning, we compared them to CSD markers of layers evoked by sensory stimulation on the mouse whisker (Supplementary Figure 2). Therefore we distinguished neurons from infragranular, granular and supragranular layers based on those boundaries.

Since intracortical column connectivity, neuronal density, membrane properties and post-synaptic targets of local PV neurons differ between various cortical layers (Hafner et al., 2019; Vecchia et al., 2020), local rhythmic PV neuronal activation should shed light on the functional implication of this varying circuit connectivity within a column. We investigated the differential impact of rhythmic entrainment of supragranular vs infragranular PV neurons across a frequency range from 0.5 to 100Hz on neuronal activity within an S1 cortical column (Figure 3a-b). Negatively-responding neurons of both supragranular and infragranular layers fired preferentially during µLED ‘OFF‘ times (Figure 3 b). However, the average phase of firing significantly varied between these infragranular and supragranular neurons (Figure 3c). Infragranular neurons fired significantly earlier after the stimulation offsets than their supragranular layer counterparts for slow-wave and δ frequency stimulations (Figure 3c; for example at 4Hz, infra-neurons:, n = 311; fire earlier than supra-neurons : n = 95; p < 0.001, Kuiper test). With increasing stimulation frequencies, this firing phase difference first reduced, and then inverted such that infragranular neurons fired significantly later than supragranular neurones at 40 Hz (n=311 and 89, p-value<0.05, Kuiper-test). Next, we observed that the coherence of firing of negatively-responding neuron was also significantly different between infra- and supra-granular neurones at most stimulation frequencies (Figure 3 d). Interestingly, the modulation index of supra-, but not infra-granular neurons modulation index had a bimodal distribution with peaks at ∼5 and ∼30 Hz, respectively (Figure 4e; Hartigan dip test, p < 0.0001). A similar analysis on granular layer neurons yielded results displaying both infragranular neuron-like behavior at lower frequencies (< 4 Hz) and supraganular neuron-like behavior at higher frequencies (> 20 Hz) (n= 80, Supplementary Figure 3).

**Figure 3:**
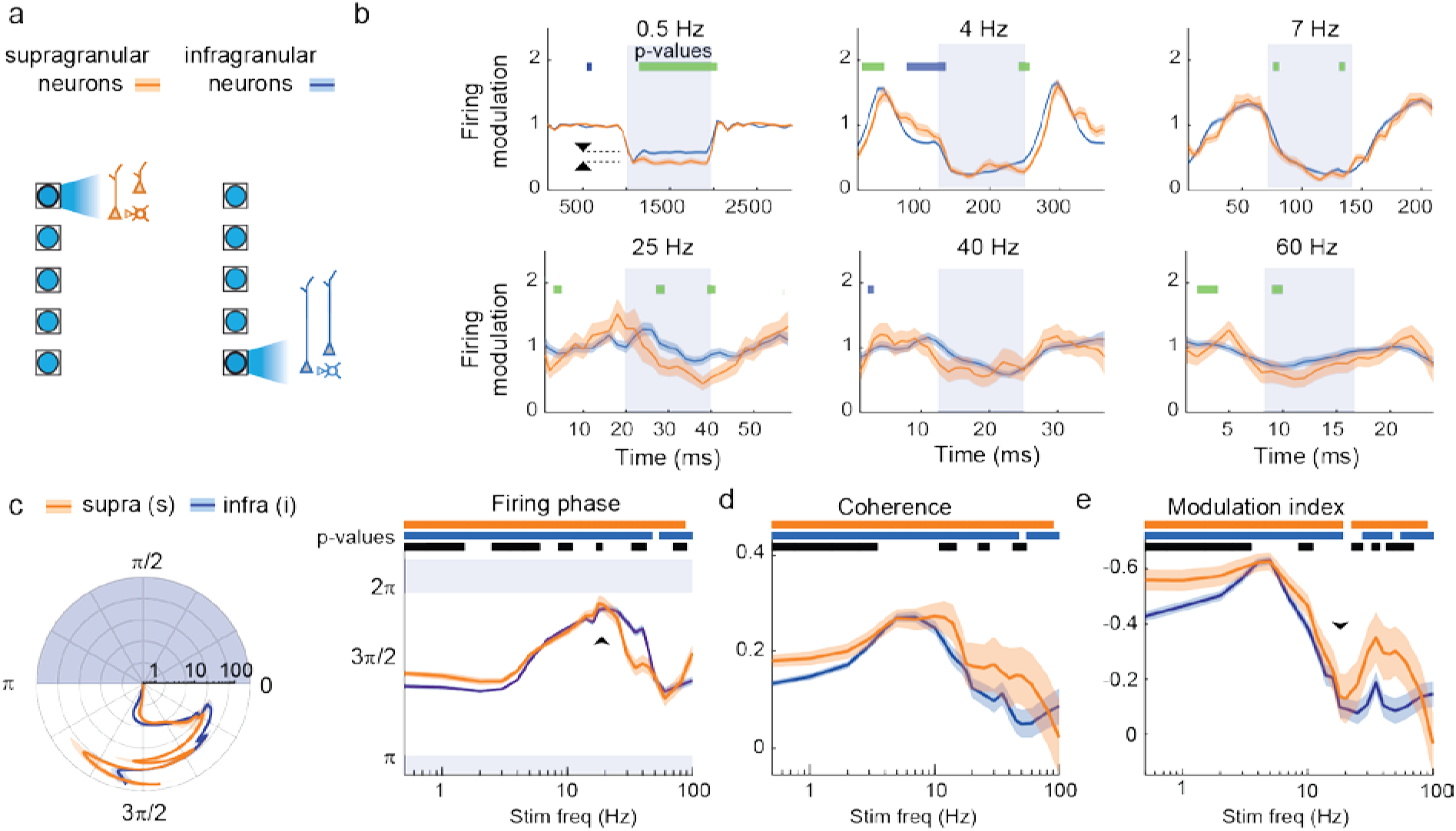
Layer-dependent entrainment of neocortical neurons. **a**) Neuronal responses of supragranular (orange), and infragranular (blue) neurons were compared when light pulse trains were delivered from local μLEDs. **b**) Firing rate responses (sliding mean ± SEM) of the neurons are represented for 6 stimulation frequencies. Significant differences at each time point were reported between the groups in blue when infragranular neurons fired less than supragranular neurons and in green in the opposite case (one-sided Wilcoxon unsigned rank test). **c**) Mean ± SEM phase of circularized firing relative to the stimulation light pulses in a polar plot (left) and orthogonal plot (right) (Blue shaded area indicates the presence of µLED stimulation). Horizontal color bars indicate significant phasing (p-value < 0.05, Rayleigh test). Black horizontal bars indicate significant phase difference between the 2 neuronal groups (p < 0.05, Kuiper test). **d**) Firing coherence across stimulation frequencies for the 2 neuronal groups. Color bars indicate p-values < 0.05, Wilcoxon one-sided signed-rank test. Black bars indicate significant difference between the 2 neuronal populations: p-val<0.05 Wilcoxon one-sided unsigned rank test. **e**) Average modulation index (mean ± SEM, note the inverted y-axis, a peak indicates a strong inhibition). The arrowhead indicates the minimum of modulation amplitude at around 20 Hz. A significant bimodal distribution was found for supragranular neurons modulation indexes (Hartigan dip test, p < 0.001).

**Figure 4.**
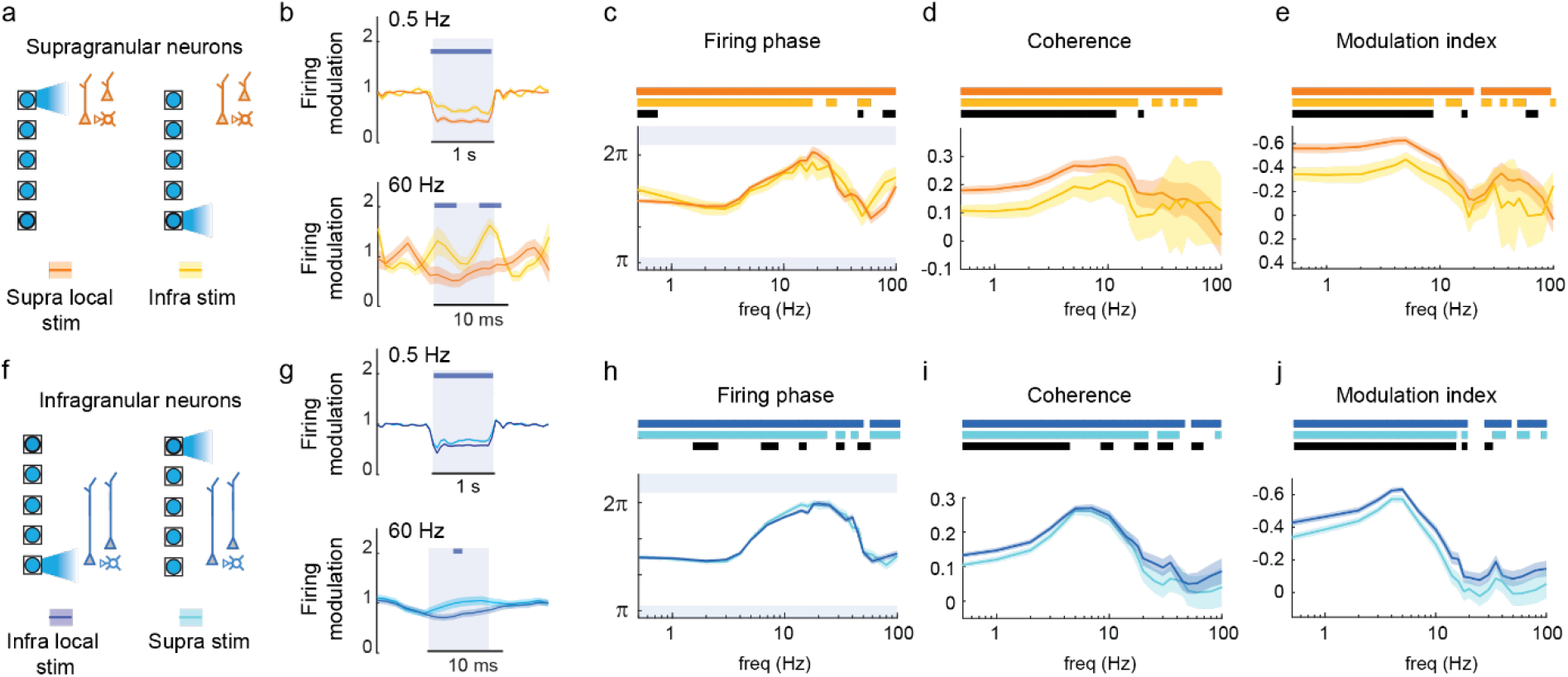
Translaminar effects of PV optogenetic activation. **a)** Schematic configuration (left) for comparing local and translaminar stimulations impact on supragranular neurons. **b**) 2 examples of firing rate responses triggered on light pulse stimulation frequency (0.5 and 60 Hz). Horizontal bars indicate significant (p < 0.05) difference between the 2 conditions for each time point. One sided Wilcoxon signed rank test (n=95 and n=84 neurons). **c**) Average angle of firing with significant difference between local and translaminar conditions (p < 0.05, Watson-Williams test black bars), and significant difference from uniform distribution (p < 0.05, Raileigh test, color bars). **d**) Coherence (mean ± SEM) of firing for the 2 conditions (bar color code : p < 0.05, Wilcoxon rank sum for 2 condition comparison, and Wilcoxon sign rank test for each separate condition). **e**) Modulation index (mean ± SEM) for the 2 conditions. Same bar code as in c for the Wilcoxon rank sum and signed rank tests. **f**) Schematic configuration (left) for comparing local and translaminar stimulations impact on infragranular neurons. **g)** Examples at 0.5 Hz (n = 368 neurons) and 60 Hz (n = 337 neurons). **h-j**) Same as (c-e) for infragranular neurons.

### Frequency-selective dampening of translaminar neuronal entrainment

After observing that cortical neurons responded in a layer-specific manner to local light stimulation, we next investigated how neuronal firing across the whole column was affected by the layer at which light stimulation occurred. Supragranular neurons (Figure 4a) were more strongly entrained by supra- vs infra-granular light stimulation (examples at 0.5 Hz (n = 95 neurons) and 60 Hz (n = 84 neurons), Figure 4b) in the slow, δ and θ frequency domains and less so in the γ frequency domain (Figure 4e). Similarly, infragranular neurons were preferentially entrained by local than by supragranular stimulations at stimulations frequencies in the slow, δ and θ ranges (Figure 4f-j, examples at 0.5 and 60 Hz in 4g for n = 368 and n = 337 neurons respectively) and to some lesser extent in the γ range as well. Overall, the inhibition generated through PV-neuron rhythmic activation spreads across layers with some attenuation particularly strong in the slow, δ and θ frequency domains. In addition, although both infra and supragranular neurons are better entrained by local than remote PV neurons stimulation, the difference was more pronounced in supragranular layers (Figure 4d-e vs. 4i-j, for example the coherence dropped by 32% from 0.25±0.02 to 0.17±0.03 for supragranular neurons whereas it dropped by 8% from 0.25±0.01 to 0.23±0.01 for infragranular layer neurons, indicating a preference for the propagation of inhibition from supragranular to infragranular layers rather than in the other direction (i.e. from infra to supra). Therefore, this questions how the neuronal specific properties across layers contribute to the propagation of inhibition.

### Differential local entrainment of excitatory and inhibitory neurons

Given their functional differences, we aimed to distinguish the respective contribution of negatively-responding INH inhibitory and negatively-responding EXC excitatory neurons to the entrained populations (Figure 5 and Supplementary Figure 4). On the basis of their spike waveforms (narrow vs broad, Supplementary Figure 1d), putative inhibitory (INH) and excitatory (EXC) neurons were separated for the estimation of firing phase, coherence and modulation index of neuronal firing as a function of local PV stimulation frequency. In supragranular layers (Figure 5a), firing phase did not differ between INH (n = 38) and EXC neurons (n = 58) (Figure 5b). The coherence and the modulation index were significantly stronger for the EXC neurons compared to INH neurons, in the slow, δ and θ frequency domains (Figure 5c, d). In contrast, in infragranular layers (Figure 5e), INH (n = 39) and EXC (n = 272) neurons showed mostly non-significantly different coherence and modulation indexes in those frequency domains (Figure 5g-h). However the firing phase was significantly different between EXC and INH in the γ frequency domain (Figure 5f). In the granular layer (Supplementary Figure 4a), EXC neurons were significantly better entrained than INH neurons in the slow, δ, θ and γ frequency domains (Supplementary Figure 4b-d). In addition, EXC but not INH supragranular neurons, showed significantly better entrainment than their infragranular counterparts in the slow, δ and θ frequency domains (Supplementary Figure 5a-b), indicating that infra- vs supra-granular differences observed in Figure 3d-e, could be driven mostly by EXC neurons, whereas the effect in the γ frequency domain could be driven by both type of neurons.

**Figure 5:**
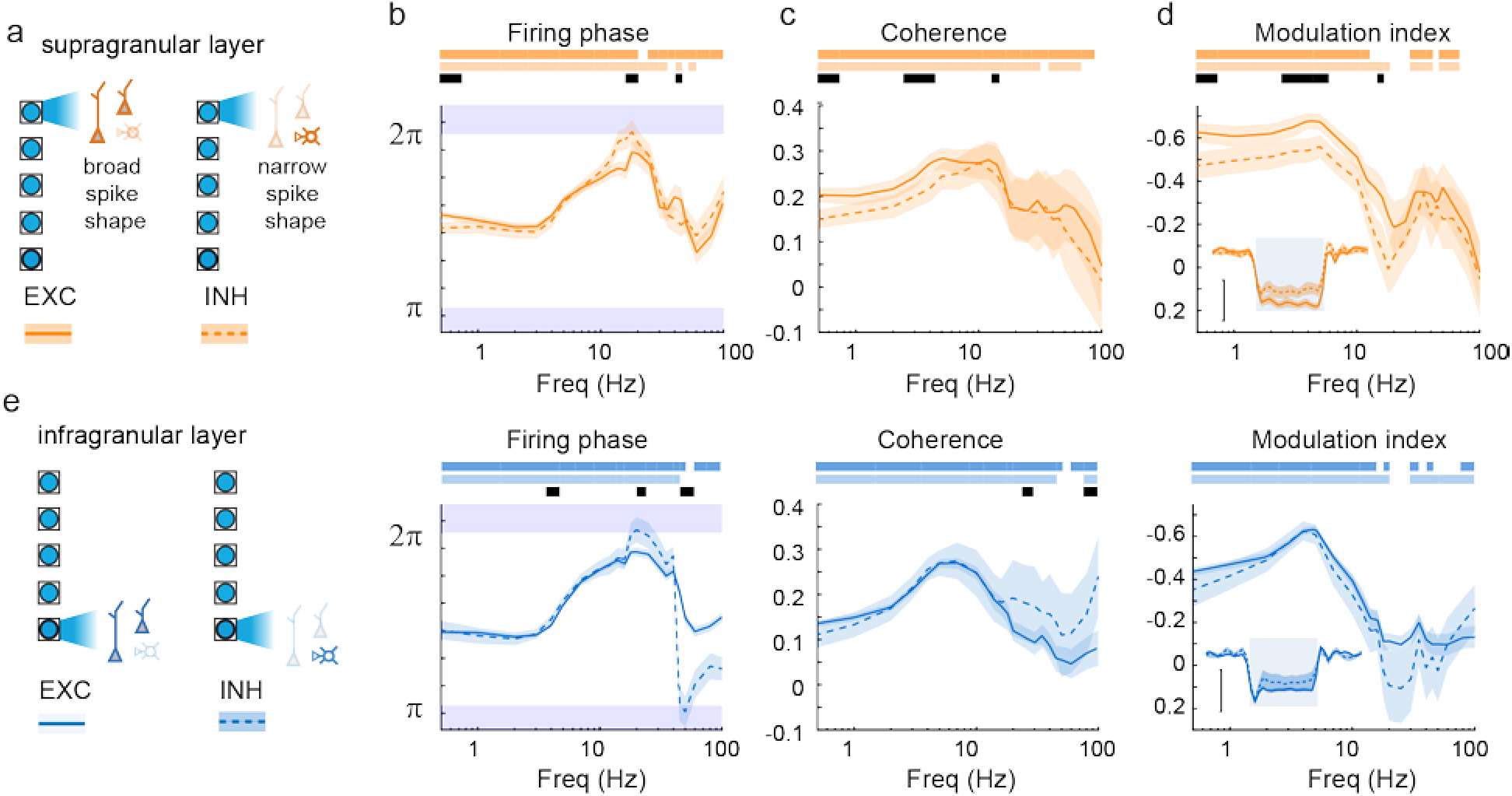
Cell type-specific local neuronal entrainment. **a-d)** Comparison of entrainment properties: firing phase, coherence and modulation index of supragranular neurons entrained by supragranular optogenetic PV stimulations, dashed lines: INH (or narrow spike shaped) vs continuous lines: EXC (or broad spike shaped). Inset represent the temporal response Vertical scale bar indicate 50% reduction of firing rate in response to 1 sec stimulation (shaded blue). **e-h)** Same as a-d for infragranular neurons with infragranular light stimulation. Color bars indicated p-values < 0.05 with phasing (Rayleigh test), coherence (Wilcoxon signed rank test) and modulation index (Wilcoxon signed rank test) respectively significantly different from uniform or null distribution respectively. Black bars indicate significant difference (p-value < 0.05) between neuronal types (Wilcoxon rank sum test).

### LFP rhythms induced by PV-neuron activation reveal complex network interactions

EEG and cortical LFP oscillatory activities are good readouts of cognitive processes (Ding et al., 2016; Bastos et al., 2020; Hayden et al., 2021) and vigilance states (Vyazovskiy et al., 2011). Therefore, we investigated how these signals were affected by our layer-specific optogenetic stimulation of PV neurons. When stimulations were applied at δ frequencies, both infra- and supra-LFP signals displayed an increase in δ-oscillation amplitude, whether or not light was applied in their respective layers (Figure 6a) indicating that a subset of cortical PV neurons is sufficient to entrain the LFP signals within the entire cortical column. However, this entrainment effect was reduced at γ frequencies (Figure 6b). To compare quantitatively the impact of these stimulations on supra- and infra-LFPs, we computed an LFP modulation index based on the amplification of local LFP measurement (Kajikawa and Schroeder, 2011, see Methods). The modulation index increased significantly at δ frequencies, for all depths of stimulations (Figure 6c, yellow bars) for supra-LFP (Figure 6c left, maximal modulation = 3.57 ± 0.53 at −700 µm, n = 24 trials, p < 0.0001, rank sum test) and infra-LFP (Figure 6c right, maximum modulation of 4.12 ± 0.18 at −900 µm, n = 28 trials, p < 0.0001, rank sum test). The infra-LFP modulation at δ frequencies was higher when the light stimulation was applied at the level of infragranular vs supragranular layers (Figure 6c right, indexes 1.81 ± 0.12 at −100 µm vs. 4.12 ± 0.18 at −900 µm, p < 0.0001, rank sum test, n_1_ = 28, n_2_ = 24). The impact of local light stimulation on local LFP exhibited a peak emerging at 5 Hz that is higher when the stimulation was delivered in infragranular layers (Figure 6e-g). Overall, at low frequencies (δ and θ not shown), both supra- and infra-LFP modulation were positively correlated with the stimulation depth (Figure 6c, linear regression: slope = −0.31, r^2^ = 0.20, p < 0.0001 for infra-LFPs and slope = −0.02, r^2^ = 0.02, p < 0.05 for supra LFPs).

**Figure 6:**
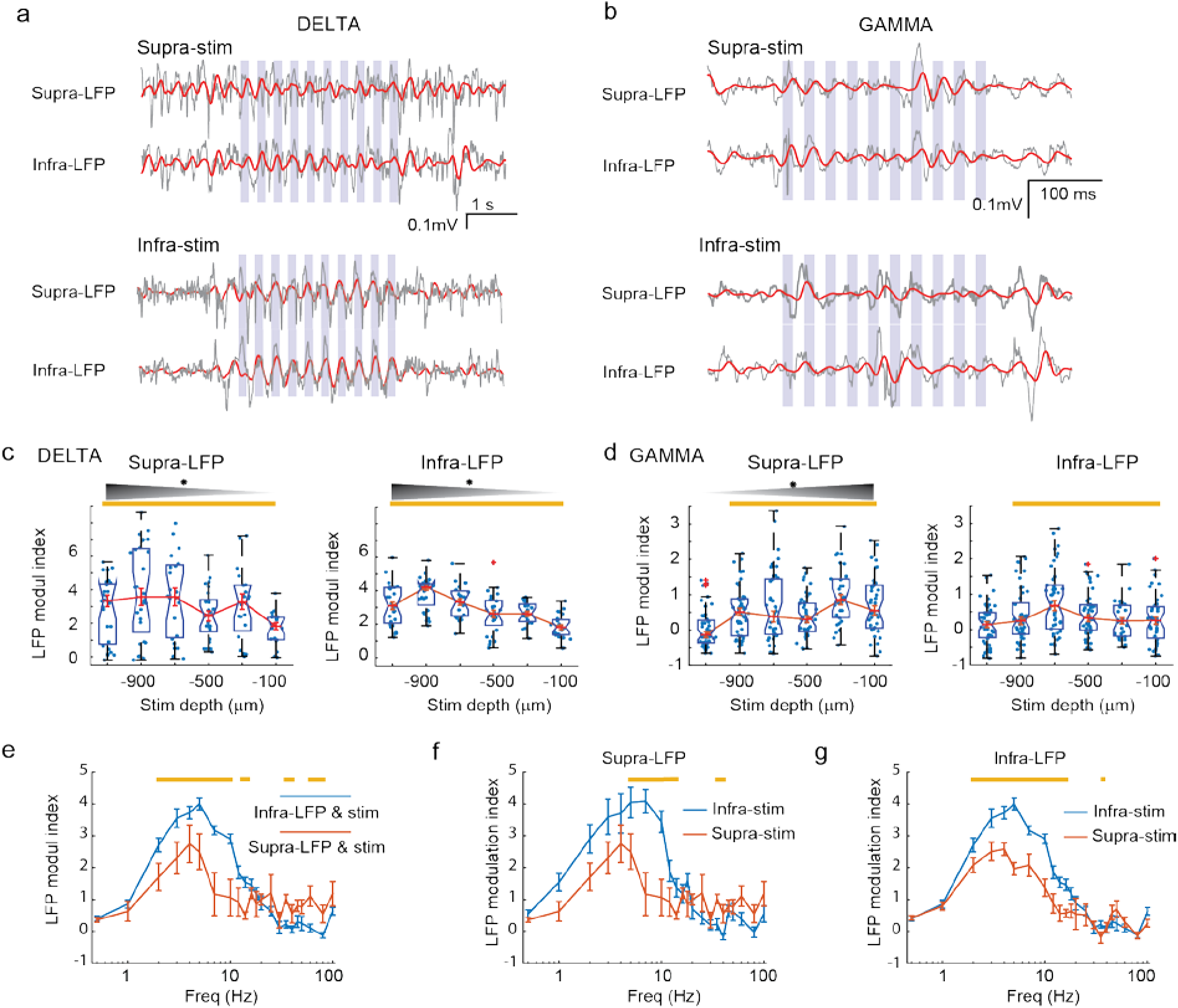
Laminar control of cortical LFP oscillations by activation of PV neurons. **a**) Example supra- and infra-granular LFP traces during 3 Hz optogenetic activation of supra- and infragranular PV neurons. Infragranular stimulations induced higher amplitude oscillations compared to supragranular layer stimulations. **b**) Same as a) for 25 Hz activation of PV neurons. **c**) Median supra- (left) and infra- (right) LFP modulation index in response to δ (2-5 Hz) frequency stimulation. Boxes indicate the 1^st^ and 3^rd^ quartiles, notches extrema and red crosses outliers. Black gradient and stars indicate significant linear correlation between LFP modulation index and stimulation depth (supra-LFP: p = 0.0074, n = 124; infra-LFP p < 0.0001, n = 128 (infragranular)). Yellow bars indicate significant modulation at each depth (p < 0.05, Wilcoxon rank sum test). **d**) Same as c) for γ (25-80 Hz) frequency stimulations. **e**) Modulation index curve of infra-LFP (blue), supra-LFP (red) in response to infra-and supragranular light stimulations. **f**) Comparison of supra-LFP modulation indexes for infra- (blue) and supra- (red) granular light stimulations. Yellow bars indicate significant differences (p < 0.05, Wilcoxon signed-rank test)) **g**) Comparison of infra-LFP modulation indexes for infra-(blue) and supra-(red) granular light stimulations.

At higher γ-frequencies, we found that local activation of PV neurons resulted in larger LFP modulation at most stimulation depths (Figure 6d, yellow bars). However, only supra-LFP modulation was linearly correlated with the stimulation depth (Figure 6d left, supra-LFP modulation index at −300 µm = 0.85 ± 0.15, p < 0.0001, n=30 trials, significantly higher than at infragranular depths of stimulation; supra-LFP modulation index at −700 µm = 0.38 ± 0.16, p = 0.031, n=41 trials; linear regression: slope = 0.12, r^2^ = 0.07, p < 0.0001), indicating that gamma LFP frequencies do not easily spread to supragranular LFP when generated in infragranular layers. The LFP modulation index in the γ frequency range at 35, 40 and 80 Hz was higher for supra-versus infra-LFP, when the stimulation was delivered in the respective layers (Figure 6e). The γ modulation indices remain nonetheless significantly lower compared to δ modulation indices (p<0.0001, at 4Hz vs 40Hz, Wilcoxon rank sum test).

Taken together, our results indicate that δ-θ LFP oscillations are particularly well entrained throughout a cortical column when the infragranular layers are stimulated, while γ rhythms are best entrained by local stimulation of supragranular PV neurons. This layer- and frequency-dependent LFP modulation (Figure 6e) can be compared to the entrainment properties of neuronal firing (Figure 3-5). Despite similar frequency-dependent aspects, supragranular firing was more modulated at slow and δ frequencies compared to infragranular layer neurons, whereas infragranular stimulation was most effective to modulate LFPs.

### CSD amplitude correlates with local neuron entrainment

Earlier we found a better entrainment of supragranular neurons in the slow and δ frequency domains (Figure 3d,e), while observing a better amplification of those rhythms when stimulating in the infragranular layers (Figure 6e-g). In order to better link neuronal firing and LFP entrainment properties, we used current source density (CSD) as a more local indicator of electric field generators. We computed the CSD across neocortical depths and evaluated the amplitude of its oscillating component within seven manually-delimited electro-anatomical compartments (Figure 7a, b; see Methods for the estimation of compartmental delimitations). CSD amplitudes were subsequently z-scored for all compartmental depths in order to compare their relative contributions. In supragranular layers (corresponding to the a-compartment), CSD modulation index was higher when the stimulation was delivered in supragranular layers (Figure 7c). Conversely, CSD modulation index was higher in infragranular layers (corresponding to g-depths) when the stimulation was delivered within infragranular layers (Figure 7g). For δ and θ frequencies, the modulation in the g compartment was stronger when the stimulation was in infragranular layers (Figure 7h,i). At γ frequencies, the CSD modulation was higher only when the stimulations were delivered within supragranular layers (Figure 7f).

**Figure 7:**
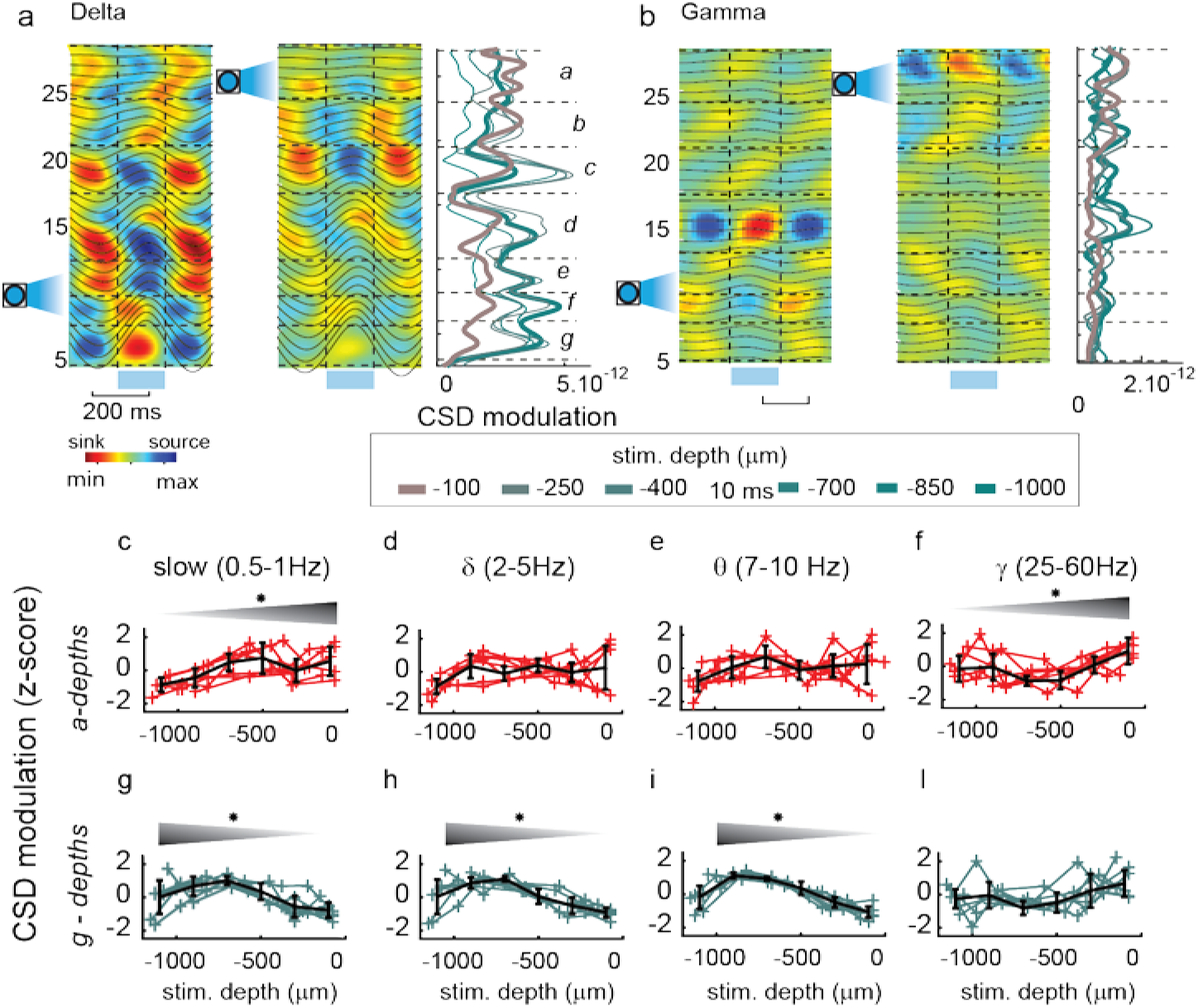
Layer- and frequency-specific current source densities during stimulations. **a**) *Left and middle panel*: Example of cortical CSD for stimulations at 3Hz for 2 μLED positions: infragranular (left) and supragranular (right) with corresponding overlaid filtered LFP traces. Shaded horizontal blue bar and vertical dashed lines indicate the μLED stimulation. *Right panel*: amplitude of the CSD map along the depth axis. The stimulation depth is color-coded. 7 depth band (*a* to *g*) are identified and delimited by horizontal dash lines. **b**) Same as in ‘a’ for a 45Hz-frequency stimulation. **c-i**) CSD amplitudes (z-scored across all stimulation depths) vs stimulation depths for n=7 mice (color) and mean±SD (black error bars) for 3 electro-anatomic compartments (i.e. *a, f*, *g)* and 4 stimulation frequencies (slow, δ, θ and γ). Top shaded triangles indicate a significant effect of stimulation depths (x-axis) either in supragranular or in infragranular layers (p-values estimated on the linear regression across stimulation depths).

The higher CSD modulation at slow frequencies (Figure 7c) and higher modulation of neuronal firing rate in supragranular layers (observed in Figure 3d,e) are now comparable, indicating a decoupling between firing response and LFP measurements.

## Discussion

We investigated the impact of rhythmic, layer-specific, optogenetic stimulation of PV neuron subpopulations at the level of neuronal firing and network -local field potential or current source density-activity within a column of the mouse somatosensory barrel neocortex in non-anesthetized awake animals. Specifically, we found that supra- and infra-granular PV populations differentially entrained columnar networks at certain rhythms and that these locally-generated entrainments could spread across layers.

In all layers, we found common frequency preferences for oscillatory entrainment in the range 2-16 Hz with a peak at 5Hz at the level of the LFP and of the neuronal firing of excitatory and inhibitory neurons. In comparison, γ-frequency stimulations entrained the LFP and neuronal overall firing rates with less amplitude. We also found that this frequency-dependent entrainment was layer-specific by using μLEDs that allowed us to optogenetically target PV neuronal subpopulations with high spatial resolution within a cortical column. When subsets of infragranular PV neurons were entrained at δ/θ frequencies, LFPs were in turn better entrained (throughout the column) than when supragranular PV were stimulated, while the opposite was true at γ frequencies (Figure 6). Consistent with this observation, firing rates of supragranular neurons were better modulated compared to infragranular neurons when either group was entrained at γ frequencies (Figure 3d,e). Similarly, supragranular layer neurons firing rates were more strongly modulated than infragranular layer neurons at slow-δ frequencies (Figure 3d,e). This layer-specific ability for supragranular neuronal firing to be well modulated at both end of the frequency spectrum was confirmed by performing CSD analysis, which indeed revealed a marked local source-sink alternation in supragranular layers (Figure 7c,f).

Lastly, EXC neurons were on average better modulated than inhibitory (INH) neurons at δ/θ stimulation frequencies in both granular and supragranular layers (Figure 5, Supplementary Figure 4), indicating that these cortical neurons are more susceptible to rhythmic entrainment.

### Conditions of layer-specific amplification of cortical rhythms

The layer-specific LFP entrainment properties that we observed in non-anesthetized mice are in agreement with previously reported layer-specific properties of cortical rhythm generation, notably that deep layer neurons are specifically capable of generating δ waves in rats (Lőrincz et al., 2015) and humans (Carracedo et al., 2013) while upper cortical layers better synchronize neuronal activity and LFPs in response to rhythmic stimulation of PV neurons at γ frequencies (Siegle et al., 2014). Our results also match with the layer-based functional organization of neocortical activity previously observed in monkeys (Buffalo et al., 2011; Bastos et al., 2020).

However, we further show that column-wide cortical rhythms could be reliably driven by a small subset of PV neurones stimulated with a single cortical layer and that these rhythms could spread across layers (Figure 4). Previous in vitro studies investigating the contribution of supra- and infra-granular layers to slow and δ frequency activity revealed the prominent role of infragranular cortex in maintaining this activity (Sanchez-Vives et al., 2000; Wester and Contreras, 2012). Similarly, slow oscillations (∼1 Hz) in mouse barrel cortex were shown to be exclusively generated within layer 5 in anesthetized animals (Beltramo et al., 2013). We found however, in non-anesthetized animals, that infra- and supra-granular layers were both efficient at entraining slow oscillations (Figure 6-7). The differences between their results and ours could be due to the anesthetized vs awake conditions. Indeed, anaesthesia has been shown via norepinephrine reduction to shunt apical dendritic communication with the soma through I_h_ activation (Phillips et al., 2016) or also to block directly amplification currents such as those emanating from T-type calcium channels activation (Timic Stamenic et al., 2019) and I_h_ inward cation non-specific current (Zhou et al., 2012). Instead, under awake, non-anesthetized conditions, inhibitory inputs within the supragranular layers could be more efficient at gating both layer 2/3 and L5 neurons through their apical dendritic arborizations that extend into layers 1-3, resulting in a widespread inhibitory impact.

Interestingly, the decrease in γ oscillation amplification, (i.e. markedly reduced for γ compared to δ-θ), could be due to the lack of post-inhibitory peaks of activity after each pulse of inhibition above 18 Hz (Figure 1g,d and 3b), thereby blocking a major boost to low frequency rhythms via resonant and amplification mechanisms. This decrease in amplitude, equivalent to a high-frequency cut-off, is often masked by the 1/frequency decrease of EEG/LFP power (see for instance ‘debiased’ LFP representations (Vinck et al., 2015) and wake power spectra (Senzai et al., 2019). This high-frequency cut-off might be more pronounced in infragranular layers compared to supragranular layers, and may subtend the layer-specific properties that we observed in the γ frequency range. Therefore, we distinguished 2 frequency ranges of interest: the δ-θ frequency band for which there was a post-inhibitory rebound and the γ frequency band for which the neuronal activity was differently controlled. Those two sets of conditions may indeed explain the differential ability of supra- and infra-granular layers in amplifying the δ-θ and the γ.

### Layer-specific δ-θ amplification mechanisms

The stronger amplification at δ-θ Hz frequencies originating from infragranular layers (Figure 6), and the stronger modulation at δ-θ Hz of supragranular layer neurons (Figure 3) appear counter-intuitive but rely on layer-specific cortical network and neuronal properties. The combination of post-inhibitory rebound (Llinás and Jahnsen, 1982) and sag currents (McCormick and Pape, 1990), are a well-known source of membrane potential resonance in both the thalamus (Leresche et al., 1991; Soltesz et al., 1991; Lüthi and McCormick, 1998; Ulrich, 2014) and the cortex (Hutcheon et al., 1996; Ulrich, 2002; Kalmbach et al., 2018). Furthermore, both T-type calcium channels and hyperpolarization activated cyclic nucleotide (HCN) channels are known to be present in pyramidal neurons of the neocortex (Talley et al., 1999; Berger et al., 2001) although they were not consistently compared across layers to our knowledge. We observed some signatures of their impact on membrane properties at the level of pulse-triggered firing rate averages, which displayed both a post-inhibitory rebound in firing (arrows of Figure 1d right, 1g) and an inhibition-induced increase in firing activity in infragranular neurons (double arrow in Figure 3b top-left). This observation could potentially be due to the different distribution of I_h_ channels observed in neocortex (Berger et al., 2001; Larkum et al., 2007; Kalmbach et al., 2018). This would also corroborate the contributing role of I_h_ in the entrainment of neocortical δ and θ rhythms (Stark et al., 2013; Schmidt et al., 2017). While this partly explains stronger infragranular layer-driven LFP amplification of δ/θ oscillations, it fails to explain the better supragranular neuronal entrainment at these frequencies. Dense lateral excitatory connectivity at the level of layer 5 between pyramidal neurons (Dantzker and Callaway, 2000; Schubert et al., 2007; Wester and Contreras, 2012; Schmidt et al., 2017) or the corticothalamic connectivity may further contribute to a higher rebound network-mediated amplification of the δ-θ cortical rhythms in infragranular layers.

In contrast, in supragranular layers, the less efficient filtering of the frequency response, the sparser lateral connectivity of layer 2-3 neurons could induce a relative failure to amplify the δ-θ rhythms at the network level but a very good capacity for neurons to be entrained. Those parameters may explain the emergence of multiple layer-dependent neocortical rhythms like hippocampal-related neocortical θ rhythms (Sirota et al., 2008), the rapid-eye-movement (REM) sleep θ rhythm, the α-rhythm(Lorincz et al., 2009), the sleep spindles, non-REM sleep thalamic-related δ rhythms or absence epilepsy (Crunelli et al., 2020) which all involve neocortical PV-neuron rhythmic activity in this frequency band and strongly depending on infragranular layers. The exact circuit, diverse and complex neuronal and synaptic mechanisms remain to be untangled (Pena and Rotstein, 2022).

### Layer-specific mechanisms of γ oscillations entrainment

Although mechanisms γ oscillations are subject to many hypotheses, they remain elusive in the awake cerebral neocortex. An interneuron-based γ oscillation could take place in supragranular layers of the visual neocortex (Perrenoud et al., 2016) but the γ generation mechanisms are also likely to be diverse across brain areas like hippocampus or neocortex as well as specific to neocortical layers and a result of their interaction (see for example (Saleem et al., 2017)). In our study, we looked at γ oscillation as a process driven by PV rhythmic stimulation as it is likely one of the phenomenon imposed by the action of the brain through cortico-cortical interaction specifically targeting PV-interneurons (Hafner et al., 2019).

We observed a significantly higher modulation of neuronal firing in supragranular layers at γ frequencies (Figure 3d,e, Supplementary Figure 5). In our preparation, responding PV-positive neurons could inhibit other FS neurons via FS-to-FS interconnections (Tamás et al., 1998; Galarreta and Hestrin, 2002) or pyramidal neurons, and influence the network dynamics at the γ frequency as previously proposed as possible mechanisms for γ oscillation synchronization (Whittington et al., 2000; Börgers and Kopell, 2003). Indeed, our observed peak for supragranular neuronal modulation at γ frequencies matches the frequency band of maximal coherence between spikes and LFP (Perrenoud et al., 2016) and the frequency band at which PV neurons stimulation triggers maximal increase in LFP power spectra (Cardin et al., 2009). This cortical network property could result from the subthreshold resonant and spiking resonant properties which are both present in cortical PV neurons (Tikidji-Hamburyan et al., 2015). Although this layer-specific property was surprising, infragranular PV-neuron intrinsic functional properties are still considerably less well characterized than supragranular neurons and were also shown to be differently connected across layers (Campagnola et al., 2022). The existence of two populations of cortical infra- and supragranular layer PV neurons could then yield in local difference of PV-driven γ rhythms and different band-pass filtering in neocortical layers. The functional properties within a cortical column and consequences are discussed below.

### Functional implications for layer-specific computations

The layer-based architecture of the neocortex is known to segregate the sensory signal based on its nature i.e. afferent, recurrent or efferent. As the sensory information flood the supragranular layers, the need to convert sensory processing into perception was proposed to take place across layers of a cortical column and to use different frequencies (Bastos 2012). Although, its dynamic processes are still debated, Bastos et al proposed that the cortical column hosts a process of information competition between prediction errors in the high frequency range and predictions in lower frequency range. One conundrum of that computation is the biological implementation. Among the proposed mechanisms, the embedding of feed-forward information (prediction errors) in supragranular layers at higher frequencies (notably γ rhythms), and of feedback information (predictions) in infragranular layers at lower frequencies (notably beta or alpha rhythms)(Bastos et al., 2020) remains to validate. In the perspective of that hypothesis, our results on specific resonance of neocortical supragranular layers at higher frequencies would facilitate such phenomena. Although we showed a resonance of infragranular layers at lower frequencies which goes in the general hypothesis, the idea of an underlying resonance at alpha (8-14Hz) frequencies in monkeys in infragranular layers, is difficult to validate, as functional frequency bands could be different in mice and brain areas (Senzai et al., 2019).

One other implication of the layer-specific properties is the possibility of establishing phase coherence across neocortical areas within layers driven by common feed-forward inhibitory inputs. This form of long range coherence is a marker of perceptual binding (Tallon-Baudry et al., 2001) which can also be subject to dysfunction in schizophrenia (Uhlhaas and Singer, 2014). This can apply when PV neurons receive inputs from the thalamus (Zucca et al., 2019) as well as from long range connections to S1 (Hafner et al., 2019). This may be a way for cortical circuits to filter top-down and bottom-up processes in specific frequency bands with only a relatively small subset of PV positive neurons as the connection are dense (Packer and Yuste, 2011) and to integrate information of superficial layers (like contextual information (Hamm et al., 2021)) across multiple areas.

## Acknowledgements

The project was funded by the European Union 7th Framework Program (FP7/2007-2013) under grant agreement n◦600925 (NeuroSeeker).

---

**Supplementary Figure 1:**
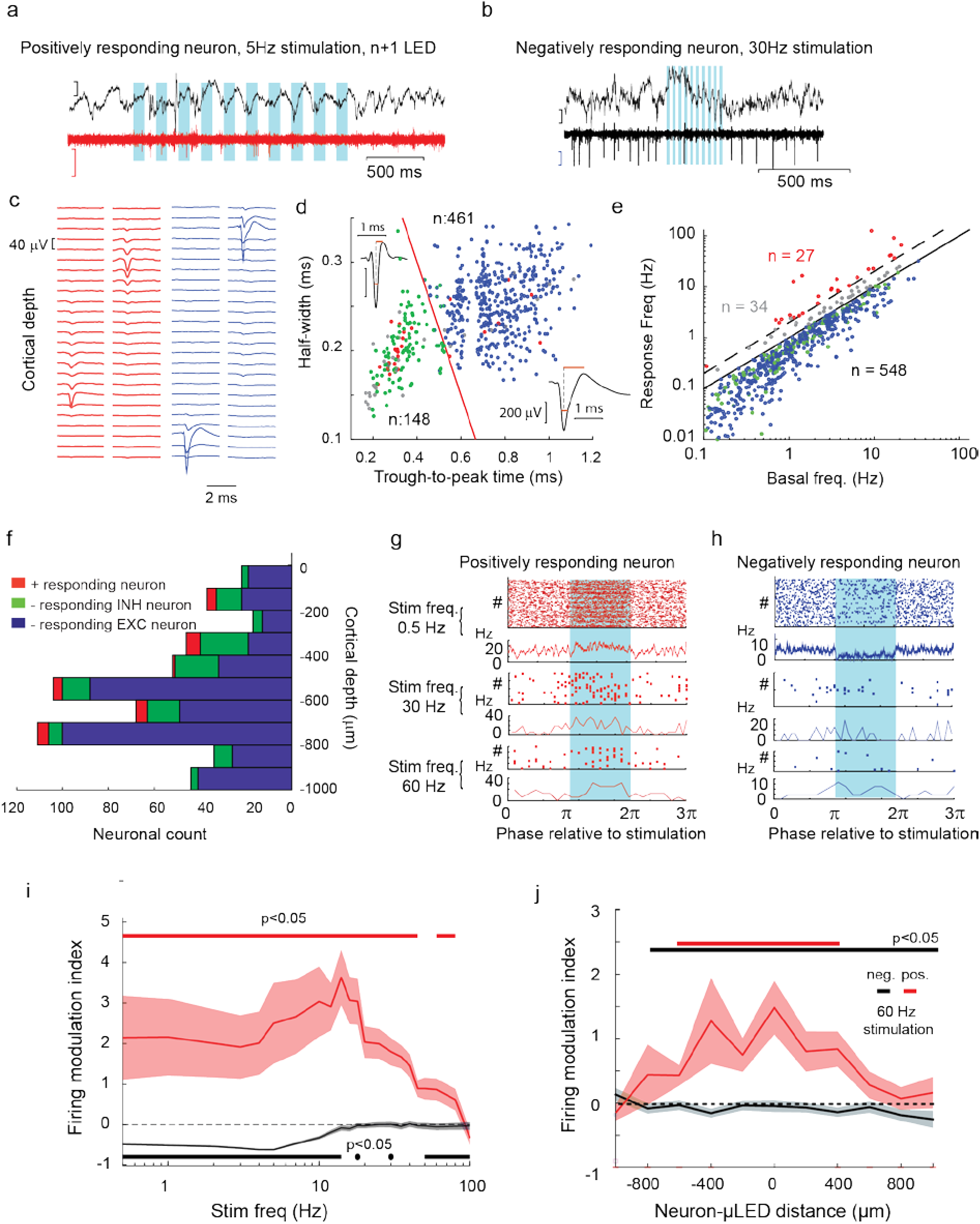
Further characterization of the neuronal responses. **a**) Same as Figure 1b left for the same positively responding unit but here for a stimulation from an µLED next to the facing µLED. **b**) Same as Figure 1b right but here for a 30Hz stimulation frequency. **c**) Spike shapes of isolated positively responding (left) and negatively responding (right) single units across electrode channels. **d**) Spike shape properties of positively-responding neurons (red) and negatively responding RS (blue) and FS (green) neurons and non-responding neurons (grey). The red line indicates the linear criterion for the classification of neuronal clusters into narrow vs. broad spike shapes. Note that the inhomogeneous repartition of blue dots is a consequence of spike shape sampling which does not affect the result. 461 clusters have broad spike shapes and 148 have narrow spike shapes. **e**) Firing frequency response (y-axis) compared to their basal firing rate (x-axis). The diagonal dashed line is the set limit between non-responding neurons (grey) and responding neurons (red), while diagonal separate negatively responding neurons from the rest. **f**) Same as Figure 1e with the EXC and INH categories. **g**) Raster plot and peri-stimulus time histogram (PSTH) for a positively-responding neuronal unit at 0.5, 30 and 60 Hz. Spike times were binned with a 1ms resolution. Blue shaded area: µLED ‘ON’ time. X-axis scale is adjusted to the period of the rhythmic stimulation **h**) Same as g for a negatively responding RS unit. **i**) Average firing modulation (± SEM) of positively (red) and negatively (blue) responding neurons to different light stimulation frequencies (x-axis). The stimulation frequencies from 0.5 to 100Hz are all represented. Horizontal bars indicate frequencies at which the modulation was significantly different from 0 (p < 0.05, Wilcoxon rank sum test). **j**) Same as Figure 1c but here for a 60Hz stimulation frequency.

**Supplementary figure 2.**
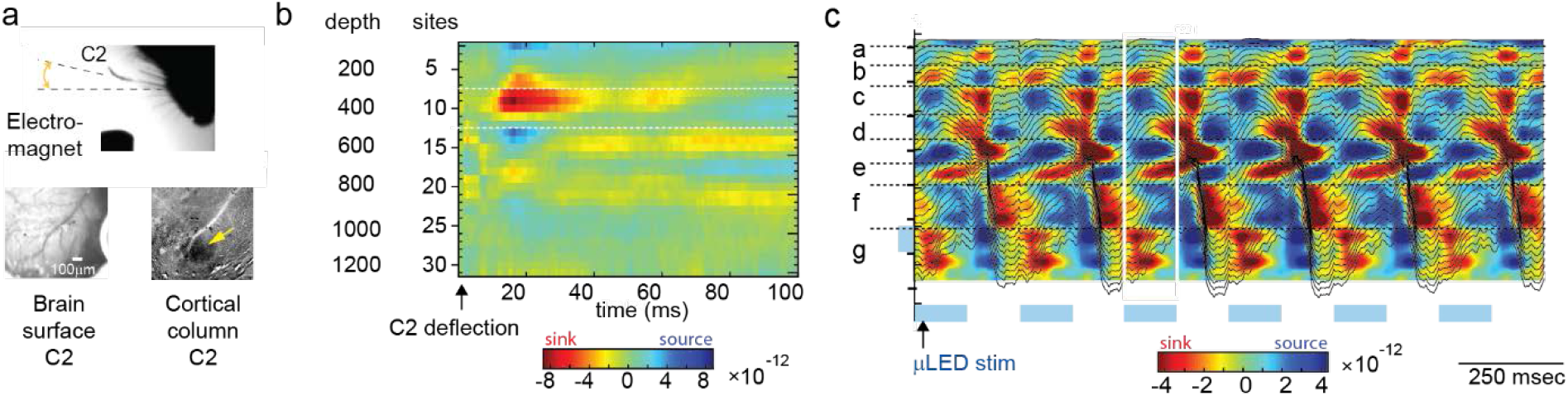
Layer specific evoked activity. **a**) Using sensory stimulations applied to the C2 whisker, the averaged CSD profile (Supplementary Figure 2b), recorded from the electrode located in the cortical barrel aligned with C2 whisker indicated a sink typically positioned across granular-supragranular positions, 20 ms following whisker sensory stimulations, as previously observed (Reyes-Puerta et al., 2015). For comparison, computing the CSD profile during the μLED stimulations, indicated a layer structured activity (Supplementary Figure 2c). **a)** Optical imaging of the barrel field (top left) and the C2 barrel column (top right) during the ferromagnetic tactile stimulation of C2 vibrissae (bottom). **b)** Current source density across cortical layers following a 5ms C2 tactile stimulation at t=0ms. The main sink 10ms after the stimulation is present in the layer 3 and 4 as previously found in the barrel aligned with the C2 stimulation (Reyes-Puerta et al., 2015). The same profile is observed in every 4 mice subject to sensory stimulation. **c)** Current source density average for 4 Hz optogenetic stimulation trains. Sinks and sources are locked to the optogenetic periodic stimulation (shaded blue bottom blocks) delivered here in the deep part of the cortex (shaded blue left block). The depth profile of distribution indicates stripes (from a to g) on which current sinks and sources alternates.

**Supplementary Figure 3:**
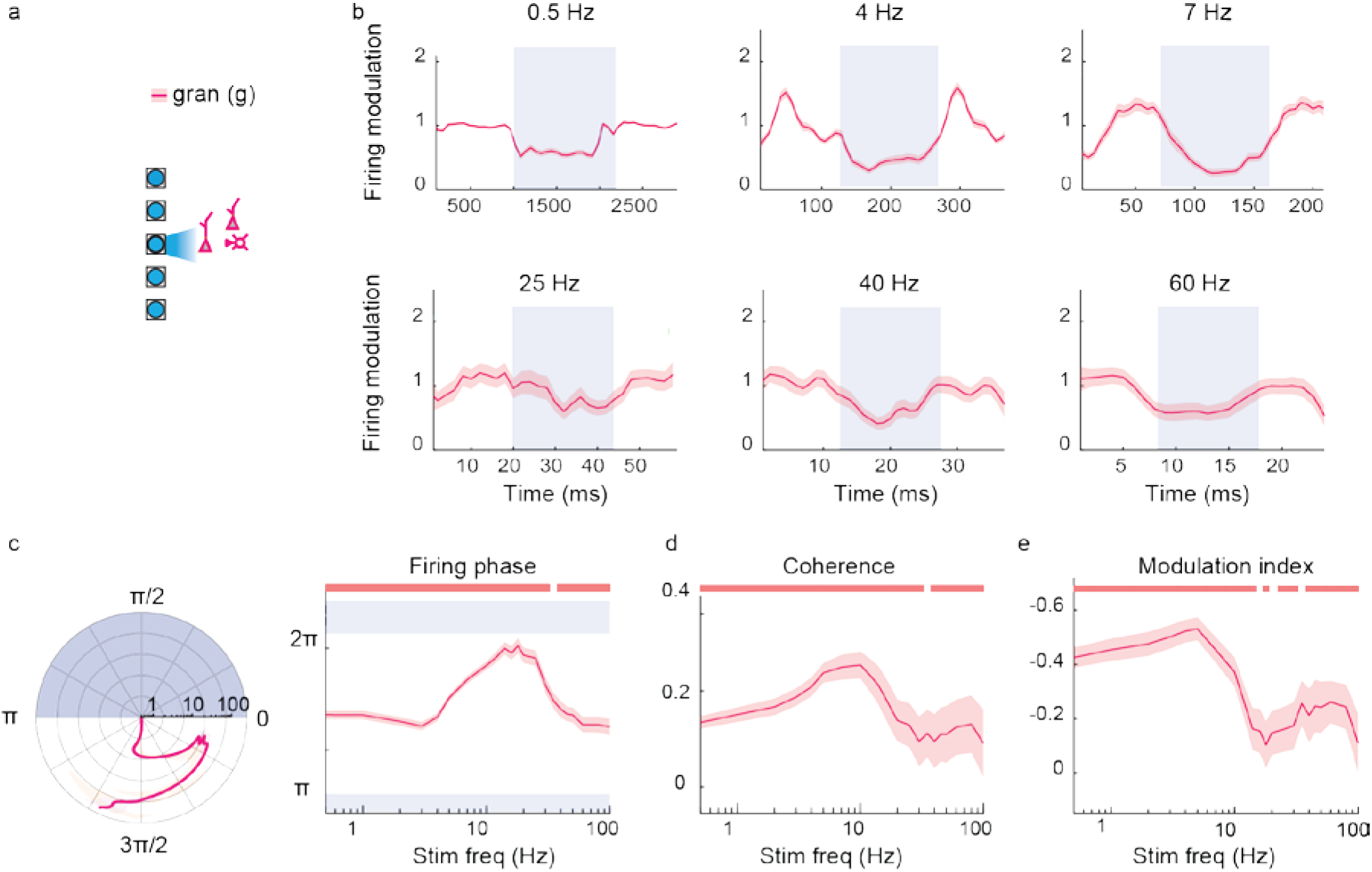
Same as figure 3 with granular layer neurons (magenta).

**Supplementary Figure 4:**
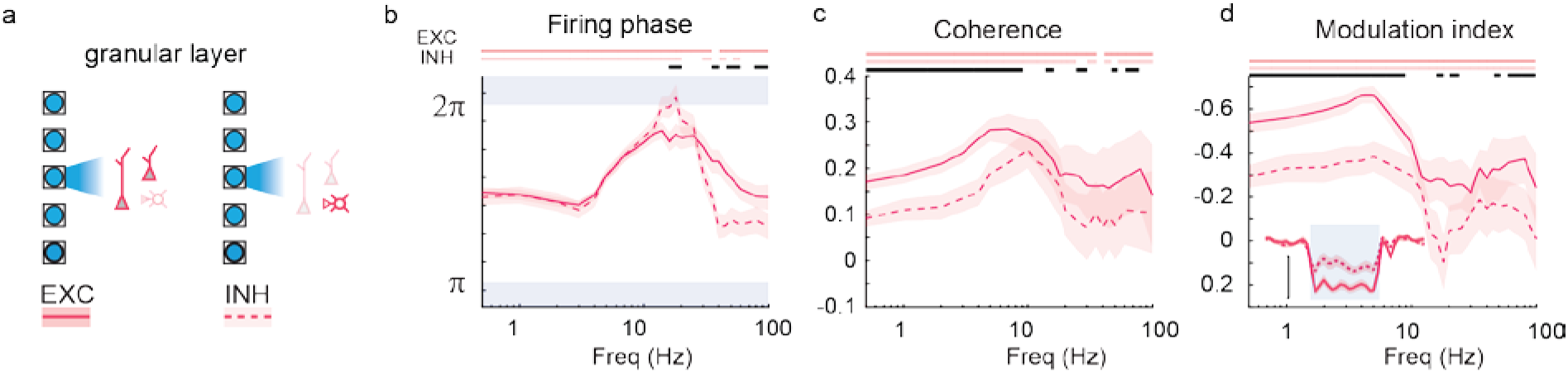
Cell type-specific local granular layer neuronal entrainment. **a-d** Entrainment analysis for the EXC and INH neurons of the granular layer.

**Supplementary Figure 5:**
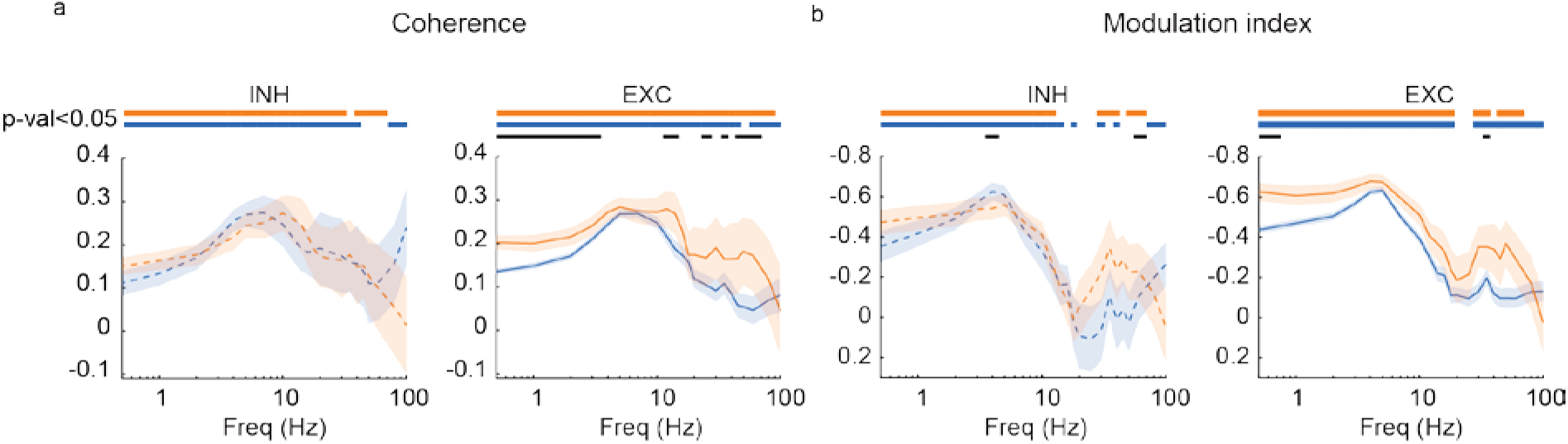
Cell type specific entrainment comparison. **a**: Comparison of coherence between supragranular and infragranular layers of INH (left) and EXC (right) neurons. p-values are indicated as follows: i/g i.e. infragranular vs supragranular layers and so on. **b**: Same as a for the modulation index here. Dash line are INH neurons, whereas plain line EXC neurons.

## Notes

### Competing Interest Statement

The authors have declared no competing interest.

### Summary of Updates

The analysis of spike was redone. The CSD analysis was added.

